# Nanolipoprotein mediated Her2 protein transfection induces malignant transformation in human breast acinar cultures

**DOI:** 10.1101/2020.08.25.265512

**Authors:** Wei He, Angela C. Evans, Matthew Coleman, Claire Robertson

## Abstract

Her2 overexpression is associated with an aggressive form of breast cancer and malignant transformation. We sought to determine if a nano-carrier system could introduce Her2 protein to non-malignant cells and determine the effects on cell phenotype and transcriptome. With stimulation with Her2-NLPs, we observed uptake of Her2 protein and a decreased probability of individual non-malignant cells forming polar growth arrested acinar-like structures. The NLP delivery system alone or Her2-NLPs plus trastuzumab showed no effect on acinar organization rate. Transcriptomics revealed essentially no effect of empty NLPs versus untreated cells whereas Her2-NLPs versus either untreated or empty NLP treated cells revealed upregulation of several factors associated with breast cancer. Pathway analysis also suggested that known nodes downstream of Her2 were activated in response to Her2-NLP treatment. This suggests that Her2-NLPs are sufficient for malignant transformation of non-malignant cells and that this system offers a new model for studying cell surface receptor signaling without genomic modification or transformation techniques.

## Introduction

Half of all drugs that have been approved to treat cancer in the past decade target cell surface receptors^1^. Despite the clinical success of these compounds, drug resistance remains an issue: for example, Her2 (Human epidermal growth factor family receptor 2, aka ERBB2) overexpressing breast cancers are treated with Her2 inhibitors such as trastuzumab, pertuzumab, and nivolumab, but ~15% of Her2 overexpressing cancers do not respond to these drugs at baseline^2^ and most patients develop resistance within 1 year^3^.

The signaling downstream of Her2 which promotes growth and malignancy is a potential source of alternative targets; however, studying this signaling is complicated by the growth advantage provided by Her2^4,5^. Some groups avoid this selection by using artificially dimerizing Her2 transgene systems^6–8^, but these systems differ significantly in their activation from wildtype Her2. Thus, we sought to determine whether a recently described protein delivery system could transport Her2 to breast cells and whether this culture system could be used as a new model of Her2-overexpressing breast cancer.

Nanolipoprotein particles (NLPs), comprised of lipid surrounded by apolipoproteins, allow for efficient solubilization of functional membrane embedded proteins^9–14^, and transfer to cells without transgenic modification or artificial cell selection^10^. NLPs, in contrast, to other protein delivery systems, do not require cell membrane disruption^15–18^ and diffuse readily due to their small size relative to silica or PLGA nanoparticles^10,13,19–21^. Furthermore, NLPs maintain correct folding of mammalian cell surface receptors^11,12,14^ whereas many delivery systems risk denaturing the proteins they carry, either by harsh packaging or endosomal lytic processes^18,22,23^. Importantly, Her2 supported NLPs retain tyrosine kinase functionality and sensitivity to Her2 inhibitors, even when synthesized in prokaryotic lysates^13^.

We chose to determine the effects of Her2-NLPs in the breast acinar morphogenesis model. Non-malignant cells cultured in 3D laminin-rich gels form growth-arrested acinar-like structures that resemble normal breast, whereas malignant cells form apolar structures growing tumor-like colonies^24,25^. Importantly, signaling downstream of Her2 has been shown to block acinar morphogenesis in similar models^6–8,26^: if Her2 transported in NLPs is sufficient for malignant transformation, cells will organize into tumor-like structures instead of acinar-like structures. We found that NLPs readily transported Her2 to non-malignant breast cells, induced tumor-like phenotypic changes and transcriptomic changes.

## Results and Discussion

### Her2-NLPs transfer functional Her2 protein to cells in 3D culture

Her2-NLPs and control empty NLPs (lipid and apolipoprotein only) were synthesized in a *E. coli* bacterial cell-free lysate and affinity purified as previously described^13^. We checked for presence of endotoxin and found that Empty NLPs contained an average endotoxin level of 104EU/mg total protein and Her2-NLPs contained 160EU/mg total protein. When diluted to 5μg/ml for cell culture, final endotoxin concentrations were less than 1 EU/ml, which represents a commonly used threshold for “endotoxin free” cell culture media. Non-malignant HMT3522-S1 cells in 3d culture were then stimulated with 5μg/ml Her2-NLP or Empty-NLPs. Non-malignant cells treated with Her2-NLPs demonstrated by positive staining for Her2 at 18 hours of culture (Supplementary Fig. 1).

### Her2-NLPs cause non-malignant S1 cells to disorganize into malignant like structures

To determine the phenotypic consequences of NLP mediated Her2 transfer, we cultured HMT-3522-S1 and -T4-2 cells in 3D laminin rich ECM (lrECM) hydrogels as previously described^25^. Briefly, single cells were dispersed in LrECM and allowed to gel, then overlaid with either culture media, or culture media +5 μg/ml of Her2 NLPs. Media and NLPs were replaced every 2-3 days for 10 days. Samples were then fixed and stained to manually identify features associated with cellular polarity. Power analysis revealed that for a chi-square test assuming an effect size of 0.33, and a significance level of 0.05, scoring 100 structures would give a power of >0.8, thus a total of 100 structures per condition per experiment were scored.

As previously reported, the majority of non-malignant S1 cells (Fig. 1A) formed growth arrested (<4% contained a mitotic figure) polar structures (60%± well organized) whereas almost all malignant T4 (Fig. 1B) cells formed apolar structures, with 30% of structures displaying 1 or more mitotic figures at the end of culture. Her2-NLP treated S1 cells (Fig. 1C) were significantly more likely to disorganize and to contain mitotic figures, compared to untreated S1 across 5 independent experiments (Pearson’s Chi-square test p<10^-14^). To ensure that the observed effects were due to Her2 and not the nanocarrier system, we repeated NLP stimulation experiments with Empty-NLPs (Fig. 1D) comprised of lipid and apolipoprotein alone at 5μg/ml, and with Her2-NLPs + a Her2 function blocking antibody inhibitor (4C5-8, a trastuzumab biosimilar) at a 1:2 molar ratio (Fig. 1E). Trastuzumab is a monoclonal antibody that is clinically used to treat HER2 positive breast cancer by binding to a conformational epitope of Her2 protein in its extracellular juxtamembrane domain and block its function. In our previous study, we have demonstrated that 4C5-8 binds to cell-free produced Her2-NLP similar to native Her2 protein^13^, and in the current experiment, we observed no effect of either empty-NLPs or Her2-NLPs in combination with Her2 targeted antibody that blocks receptor function (Fig. 1F). This shows that Her2 itself drives nonmalignant cells to disorganize and fail to growth arrest, while the supporting NLP system has minimal phenotypic effects. Titrating the Her2-NLP concentration revealed dose response behavior, with increasing probability of disorganization with increasing levels of delivered Her2 (Fig. 1G). Increased numbers of mitotic cells were also observed in Her2-NLP treated structures relative to either non-malignant or empty-NLP treated cells (Fig. 1H).

**Figure 1:**
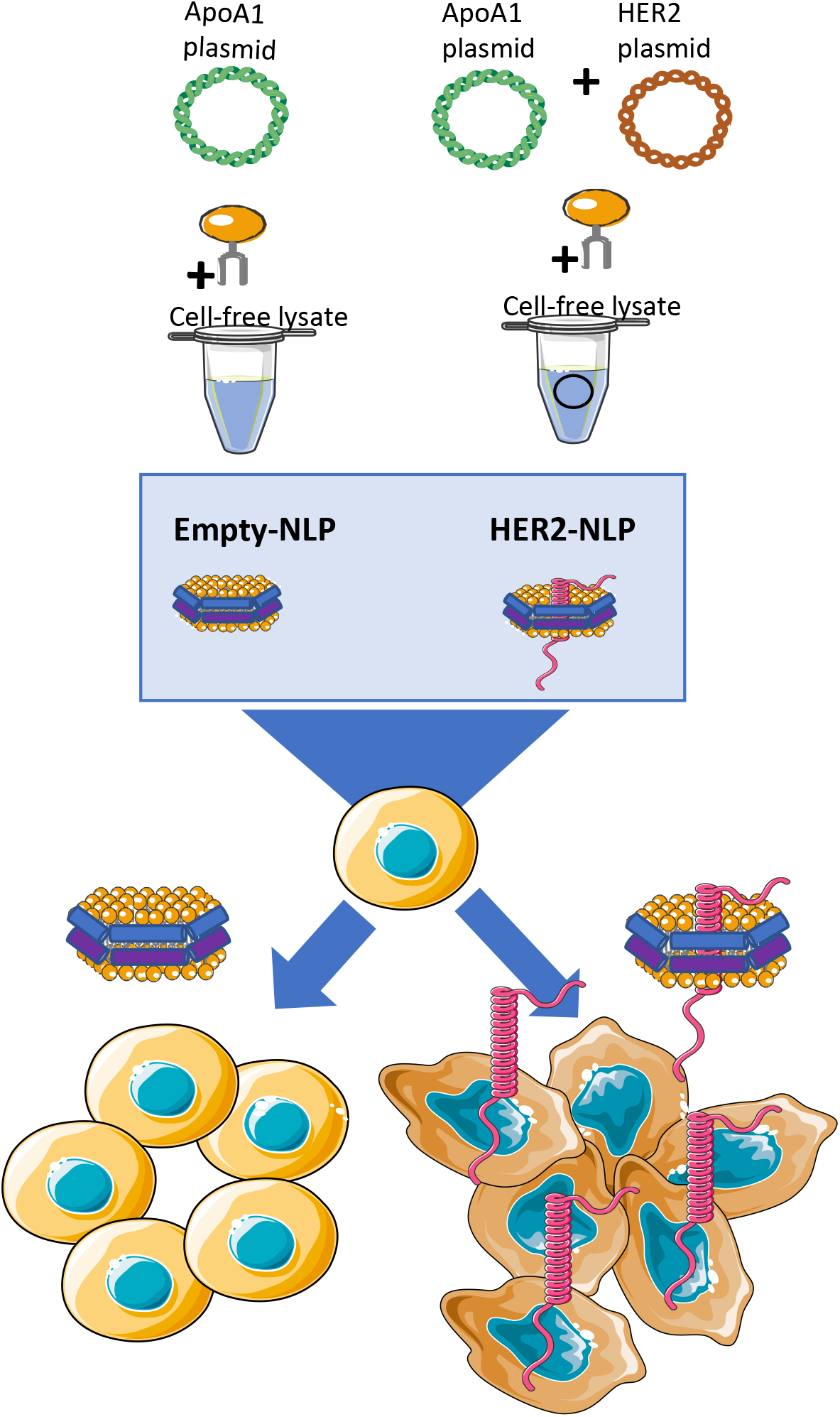
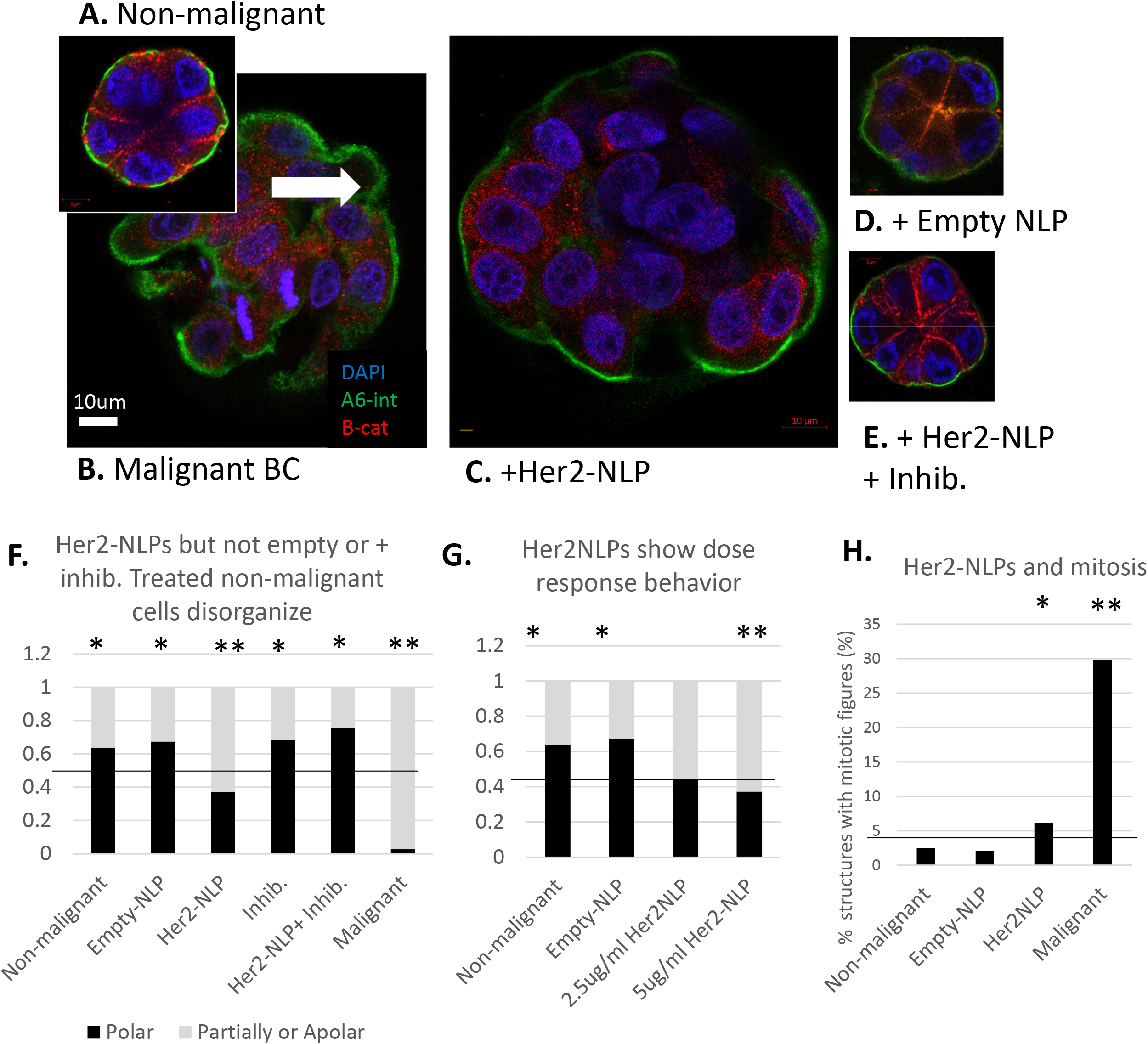
Her2 carried by NLPs causes malignant-like organization in non-malignant cells. A: Non-malignant cells cultured in lrECM form growth arrested polarized acinar like structure with basal organization of α6 integrin (green), and lateral organization of β-catenin (red). B: Malignant breast cancer cells form disorganized structures with no cell polarity which fail to growth arrest as shown by presence of mitotic figures (white arrow). C: Non-malignant cells treated with Her2-NLPs form apolar masses which fail to growth arrest. D: In contrast, non-malignant cells treated with empty-NLPs organize normally. E: non-malignant cells treated Her2-NLPs and a Her2 dimerization inhibitory antibody organize normally. F: Significantly more structures organize well in Non-malignant, empty-NLP treated, inhibitory antibody only and Her2-NLP+ inhibitory antibody conditions, whereas Her2-NLP treated and malignant cells are less likely to organize into polarized, growth arrested acini (n=5 biological replicates, * indicates post hoc test significance of p<0.05, ** indicates post hoc test of p<0.001). F: Her2-NLPs show dose response behavior, with fewer structures organizing well with increasing NLP dosage (n=3 biological replicates, * indicates post hoc test significance of p<0.05, ** indicates post hoc test of p<0.001). G: Her2-NLP treated structures are more likely to contain a mitotic figure than untreated or empty-NLP treated cells (n=3 biological replicates, * indicates post hoc test significance of p<0.05, ** indicates post hoc test of p<0.001).

To the best of our knowledge, this is the first demonstration that protein transfer of an oncoprotein is sufficient to drive malignant transformation of non-malignant breast cells in 3D culture. Akin to previous work using artificially dimerizing Her2^7^, our work demonstrates that Her2 can block acinar morphogenesis, however, instead of these relatively complex transgenic strategies, we use wildtype Her2 protein. This work is distinct from previous experiments studying the effects of Her2 using genomic modification, as we do not induce genomic breaks, nor induce an immune response to cytoplasmic DNA^27^ nor select cells. We did not directly measure dimerization state in these models, however treatment with a Her2 dimerization inhibitor blocked the effect of Her2-NLPs, indicating correct folding and presentation of the receptor.

### Her2-NLPs cause transcriptomic changes in non-malignant cells

To understand the transcriptomic changes induced by increased levels of Her2, we cultured non-malignant S1 cells with no treatment, treated with 5 μg/ml empty-NLP, or treated with 5μg/ml Her2-NLP in 3D LrECM for 8-10 days. Cells were then extracted from the lrECM hydrogen, then lysed to extract total RNA for 2 biologically independent experiments. The RNA was isolated and enriched through poly-A selection and sequenced using nextgen sequencing (HiSeq, illumina). Reads were normalized, then mapped to the human genome and compared across groups using DESeq2 using a p-value cutoff of <0.05, and an absolute log_2_ fold change of >1. PCA analysis revealed that replicate was the primary cause of variability between experiments, followed by Her2-NLP treatment (Fig. 2A).

**Figure 2:**
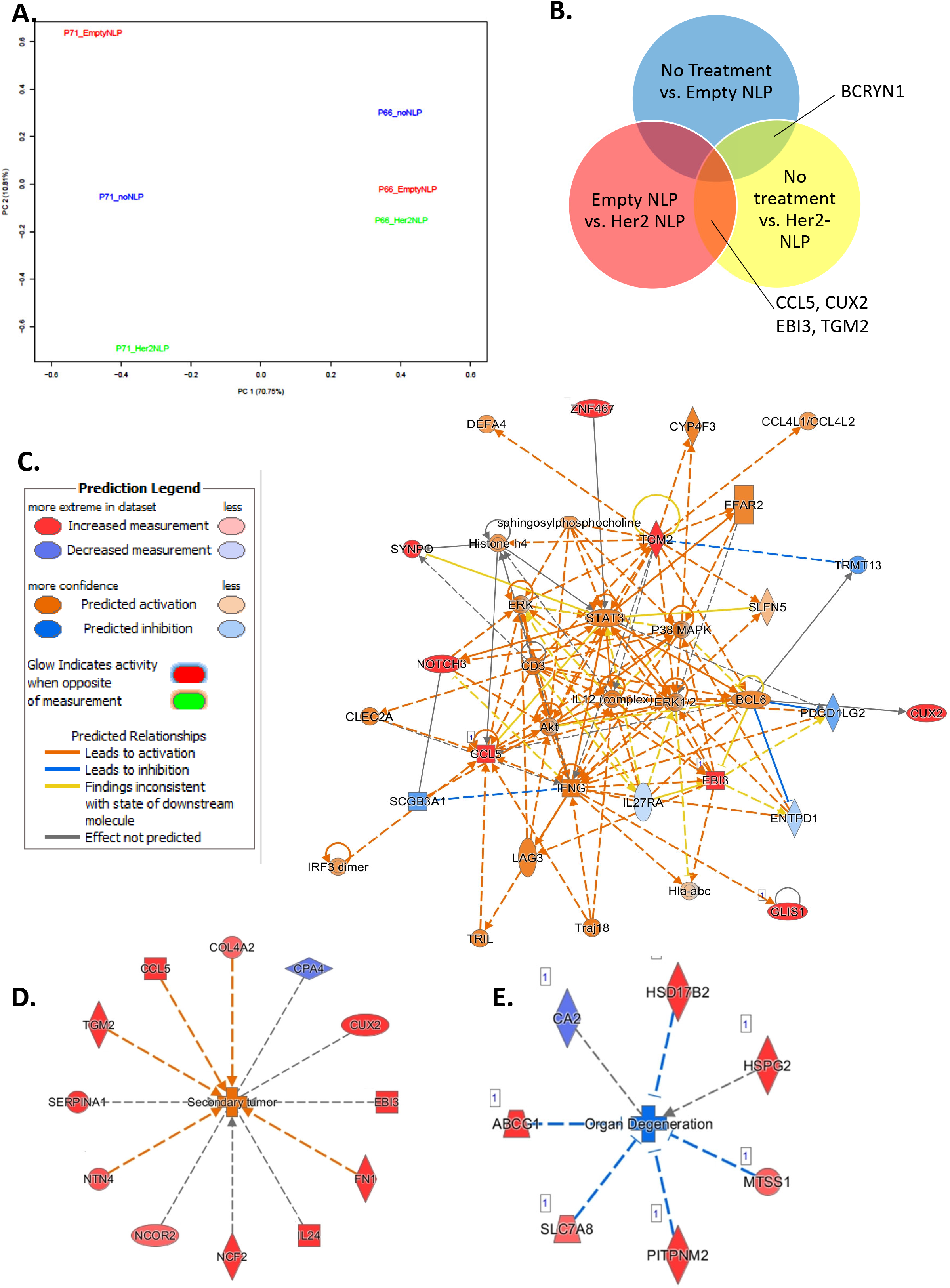
Her2-NLPs induce gene expression changes in non-malignant 3D cultures. A: PCA plot for all samples sequence shows that principle component 1 separates biological replicates, and principle component 2 separates Her2-NLP treated from other conditions. B: Genes common to multiple comparisons include BCRYN1 for all NLP treated cells versus untreated samples, and CCL5, CUX2, EBI3, and TGM2 for Her2-NLP treated samples versus either empty-NLP or untreated samples. C: Ingenuity Pathway Analysis network comparing Her2-NLPs to empty NLP discs predicts relationships among several biological families and transcriptional regulators involved in cancer progression and metastasis, including ERK, Akt, P38 MAPK, and STAT3. Genes in red indicate hits found in dataset. Orange coloring indicates predicted activation of biomolecules, and blue indicates predicted inhibition. Deeper color saturation indicates more confidence in predicted regulation. D: Secondary tumor formation is the top activated disease network associated with Her2-NLP treatment compared to untreated samples. E) Organ degeneration is predicted to be inhibited with Her2-NLP treatment compared to untreated samples. Genes in red indicate up-regulated genes found in dataset, whereas genes in purple indicate those that were down-regulated in the dataset. Orange coloring indicates predicted activation and blue indicates predicted inhibition of the biological phenotype. Log_2_ fold change+/-1, p<0.05.

Comparing untreated and empty-NLP treated cells revealed only 1 differentially expressed transcript, the noncoding lncRNA BCYRN1. Gene ontology analysis also revealed no differentially expressed terms, further emphasizing the minimal effects of the NLP system (Fig. 2B and Supplementary Fig. 2). In contrast, comparing empty-NLPs and Her2-NLPs revealed 9 upregulated genes and comparing no treatment and Her2-NLPs revealed 32 upregulated genes and 6 downregulated genes. Four genes were consistently upregulated with Her2-NLP treatment as compared to either empty-NLP alone or untreated cells (Fig. 2B). Transcripts identified in Her2-NLP treated groups relative to either untreated or empty-NLP treated factors overlapped with previously reported screens for acinar morphogenesis (EBI3, CCL5)^28^ and for residual disease in Her2+ breast cancer (CCL5^29,30^), breast cancer outcome (EBI3^31–33^) and metastasis (TGM2^34^, EBI3^33,35^).

Differentially expressed genes comparing Her2-NLP treatment to empty-NLPs or to untreated cell cultures were analyzed via Ingenuity Pathway Analysis (IPA). Her2-NLP treated cells compared to either untreated or empty-NLP treated cells showed a clear activation of cancer-related biological responses (Supplementary Table 2) and predicted activation of signaling nodes which have been linked to Her2 overexpression such as Stat3^36^, ERK and TgFB^26^, p38MAPK^37^, and NFkB^28^ (Fig. 2C and Supplementary Fig. 3). Notably, signatures of cancer-related malignancies, including breast and ovarian cancers, were identified as prominently associated with HER-2 NLP treatment as compared to empty-NLPs (Supplementary Fig 4). The strongest disease networks associated with Her2-NLP treatment were activation of secondary tumor formation and suppression of organ degeneration (Fig. 2D-E). Her2-NLP treatment was also linked with several cancer-related disease behaviors, including proliferation, the synthesis of reactive oxygen species, and formation of cellular protrusions. Upstream factor analysis suggested that common upstream nodes may include NFκB, lipopolysaccharide, interferon gamma, and TNF (Supplementary Table 3).

Our demonstration that Her2-NLPs induce malignant like phenotypic and transcriptomic changes shows that this model represents an oncoprotein driven model of malignant transformation. Specifically, predicted activation of breast cancer associated signaling nodes suggests that our model recapitulates features of Her2 overexpressing breast cancer and may offer new avenues to study Her2 signaling in heterogeneous backgrounds. Future work using this system can take advantage of the lack of alternative splicing of Her2, dynamic introduction of Her2 for pulse-chase experiments, and use the reversible nature of this system to study Her2 withdrawal.

## Conclusions

In this work, we demonstrate, for the first time to our knowledge, that receptor-driven oncoprotein transfection can induce malignant transformation in phenotypic cultures. Specifically, we demonstrate that the Her2 receptor transferred to non-malignant breast cells induces malignant like growth patterns in a subset of cells, and that this effect is not seen either with the protein transfer system (NLP alone) or Her2-NLP in concert with a conformational dimerization inhibitor. This protein delivery system is limited to membrane bound proteins, but given the centrality of the surface receptors in cancer, including the tyrosine kinase receptor family and the G protein-coupled receptor family, there is a clear use for this system in rapidly engineering receptor driven cancers, which represent targets for over half of all cancer drugs approved in the last decade.

## Acknowledgements

Funding: Lawrence Livermore National Lab Laboratory Directed Research Program 18-ERD-062 (to CR) and 19-S1-003.

Thanks to Christine Ichim, Monica Moya, W. Rick Hynes for their helpful discussion related to this work. Thanks to Aimy Sebastian for help with bioinformatics analysis.

This work was performed under the auspices of the U.S. Department of Energy by Lawrence Livermore National Laboratory under Contract DE-AC52-07NA27344. IM release number LLNL-JRNL-813643

## Supplementary Materials and Methods

### NLP Preparation

Her2-NLPs were prepared as previously described^13^. Briefly, plasmids encoding for human full-length ErbB2 gene and a truncated 6x-His-tagged apolipoprotein A1 (Δ49A1) were synthesized using the cell-free Expressway system (Life Technologies), in the presence of 1,2-dimyristoyl-sn-glycero-3-phosphocholine (DMPC, Avanti). After overnight expression, Her2-NLPs were harvested from the cell-free mixture by native nickel pulldown^13^. The purified NLP stocks then underwent buffer exchange into pH7.4 PBS and sterile filtration for use in antibiotic free mammalian cell cultures. Empty NLPs were made using same cell-free method except for without ErbB2 plasmid. Protein concentrations were determined by Nandrop. Endotoxin concentrations were determined using the Endosafe□-PTSTM (Charles River) endotoxin testing system based on the Limulus amebocyte lysate assay and is expressed as EU/mg total protein.

### Cell Culture

Human immortalized breast cells (S1) and breast cancer cells (T4-2) from the HMT3522 progression series (kind gift of Mina Bissell)^38^ were maintained in DMEM/F12 (11330, ThermoFisher) supplemented with 250ng/ml insulin (I6634, Sigma Aldrich), 10μg/ml Apo-transferrin (T2252, Sigma Aldrich), 2.6ng/ml sodium selenite (Corning, 47743-618), 10^-10^ M beta-estradiol (E2785, Sigma Aldrich), 1.4×10^-6^ M hydrocortisone (H0888, Sigma Aldrich), 5μg/ml ovine prolactin (LA Biomedical Research Institute), and for S1 only, 10ng/ml EGF (11376454001, Roche). S1 were seeded in uncoated T75 flasks (corning) at 2e^4^/cm^2^, refed every 2 days, and passed every 7 days and T4-2 were seeded in collagen coated flasks (corning) at 1e^4^/cm^2^, refed every 2 days and passed every 5 and 3 days respectively. Cells were kept in humidified incubators at 37C and 5% CO2 supplementation, and CO2 calibrations were performed biweekly with a test kit (Fyrite, Bacharach). Cells were screened for mycoplasma contamination every 2 months (Mycoalert, Lonza).

### Acinar Morphogenesis Assay

S1 or T4-2 cells were passaged, counted, and resuspended in 100% lrECM (354230, growth factor reduced Matrigel, Corning) at 800k cells/ml or 600k/ml respectively on ice. Cells in lrECM were transferred to dishes, allowed to gel for 20 minutes at 37C, then covered with complete culture media ± 5μg/ml Her2-NLPs or empty NLPs or Her2-NLPs plus a Her2 blocking antibody (clone 4D5, MCA6092, Biorad) at 26.67μg/ml (which represented 1:1 molar ratio with Her2-NLPs). Media was replaced every 2-3 days for 10 days.

At 10 days, cultures were harvested for imaging by removing culture media and smearing cell laden lrECM onto coated glass slides (Superfrost plus, Thermo). Slides were then immediately fixed in 10% formalin for 15 minutes, washed 3x in PBS, blocked in for 1hr at room temp in IF buffer comprised of 3%BSA and 0.5% Triton X-100 in PBS supplemented with 10% normal goat serum and goat anti-mouse Fab fragment (115-007-003, Jackson Immunoresearch) to block mouse antibodies present in lrECM. Primary antibodies for beta-catenin (Ab32572 Abcam), alpha-6 integrin (555734, BD Biosciences) and laminin (L9393, Millipore Sigma) were diluted 1:100 in IF buffer and stained for 2hrs at room temperature. Slides were then washed 3x in IF buffer, and secondary antibodies (A21429, A-11006, A32723 as appropriate, Thermo Fisher) were applied at 1:500 dilution in IF buffer for 1hr at room temp followed by 3 washes with IF buffer, and 3 washes with PBS. Slides were then stained with dapi at 0.1 μg/ml for 5 minutes and mounted (Prolong Gold, Thermo Fisher) and cover slipped.

Slides were imaged on a laser scanning confocal microscope (LSM700, Zeiss) equipped with an Acroplan 40x/1.1NA water immersion lens. 100 structures per experiment were scored by a trained observer according to criteria in Supplementary Table 1. Pearson’s Chi-square test and Bonferroni post hoc tests were performed in R.

### RNAseq

After 10 days of culture, cells were harvested from lrECM by incubation with 5mM EDTA in PBS on ice for 30 minutes with gentle agitation and spun down to collect structures. Cell pellets were then snap frozen on dry ice for shipping, followed by RNA extraction, poly-A capture, cDNA synthesis, end repair and adaptor ligation. Samples were then sequenced (HiSeq, Illumina). Reads were trimmed to remove adapter sequences and poor-quality nucleotides, then aligned to the Homo Sapiens reference genome (GRCh38). Differentially expressed genes were compared using DEGseq2. The Wald test was used to generate p-values and log2 fold changes. Genes with an unadjusted p-value < 0.05 and absolute log2 fold change > 1 were called as differentially expressed genes for each comparison. A gene ontology analysis was performed on the statistically significant set of genes by implementing the software GeneSCF v.l.l-p2. The goa_human GO list was used to cluster the set of genes based on their biological processes and determine their statistical significance. To estimate the expression levels of alternatively spliced transcripts, the splice variant hit counts were extracted from the RNA-seq reads mapped to the genome. Differentially spliced genes were identified for groups with more than one sample by testing for significant differences in read counts on exons (and junctions) of the genes using DEXSeq.

### Ingenuity Pathway Analysis (IPA)

Differentially expressed genes from RNAseq studies were run through IPA (Qiagen). In total, three core analyses were run; Empty-NLP vs. No treatment, Her2-NLP vs. No treatment, and Her2-NLP vs. empty-NLP. Both direct and indirect relationships were considered for IPA mapping and statistical analysis. Log2 fold change cutoffs of+/-1 and p-values <0.05 were included in the analysis.

---

**Supplementary Figure 1: Her2-NLPs are taken up by cells.** A&B: An untreated cell cultured in 3d lrECM does not show any staining for Her2, whereas C&D: a treated cell shows abundant Her2 staining throughout the membrane and cytoplasm, but not nucleus.

**Supplementary Figure 2: Gene expression changes associated empty or Her2-NLP treatment.** A: Table of differentially expressed genes across all comparisons shows 1 DEG for no-treatment vs. empty NLP treatment. B: List of differentially expressed genes for all comparisons studied. Genes highlighted in yellow are common to both Her2-NLP treated comparisons and in green are common to either NLP treated condition.

**Supplementary Figure 3: Her2-NLP treatment activates the regulation of several key biological pathways when compared to untreated cells.** Networks identified via Ingenuity Pathways Analysis (IPA) show connections between differentially expressed genes in our dataset and cancer progression and immune regulated biomarkers, such as A: RAS and TNF, B: ERK, TGFB and PI3K, and C: ERK, p38MAPk, and NFkB. Genes in red indicate upregulated genes found in dataset, whereas genes in purple indicate those that were down-regulated in the dataset. Molecules in gray indicate associated connections as predicted through IPA. Log2 fold change cutoff+/-1, p<0.05.

**Supplementary Figure 4: Her2-NLPs induce differentially expressed genes that associate with cancer.** A: Her2-NLPs link with several cancer-related diseases and biofunctions. B: Eight differentially expressed genes in the Her2-NLP vs. empty-NLP comparison demonstrate significant contribution to cancer-related diseases and biofunctions. NOTCH3 and TGM2 were present in over 40% of the predicted cancer-related diseases, with NOTCH3 involved in 47% and TGM2 involved in 76% of hits, respectively. Log_2_ fold change+/-1, p<0.05. C: Full list of cancer-related malignancies and corresponding molecular hits when comparing Her2-NLP to empty-NLP discs.

**Supplementary Table 1:** Acinar Scoring Criteria.

| Acinar Scoring Criteria | Polar (all) | Somewhat Polar or Apolar (Any) |
| --- | --- | --- |
| Size | Compact | >20 cells |
| Mitotic spindles | Absent | Present |
| Positioning of ECM receptors | Basal | Lateral or apical |
| Cell-cell junctions | Well organized | Not organized |
| Colony Border | Smooth | Not Smooth |

**Supplementary Table 2:** Top diseases and biofunctions predicted to be activated (Z-score>2) or inhibited (Z-score<-2) with Her2-NLP treatment compared to untreated cells.

| Biological Effect | p-value | Activation z-score | Molecules |
| --- | --- | --- | --- |
| Secondary tumor | 2.91E-06 | 2.186 | CCL5, COL4A2, CPA4, CUX2, EBI3, FN1, IL24, NCF2, NCOR2, NTN4, SERPINA1, TGM2 |
| Synthesis of reactive oxygen species | 9.96E-05 | 2.382 | CCL5, FN1, IL24, IL32, NCF2, SERPINA1, TGM2 |
| Production of reactive oxygen species | 0.000156 | 2.189 | CCL5, FN1, IL24, IL32, NCF2, TGM2 |
| Organ Degeneration | 0.000238 | -2.219 | ABCG1, CA2, HSD17B2, HSPG2, MTSS1, PITPNM2, SLC7A8 |
| Proliferation of blood cells | 0.000823 | 2.426 | ABCG1, CCL5, EBI3, FN1, IL24, IL32, NCOR2, SERPINA1 |
| Proliferation of lymphocytes | 0.00144 | 2.23 | ABCG1, CCL5, EBI3, FN1, IL24, NCOR2, SERPINA1 |
| Formation of cellular protrusions | 0.00692 | 2.429 | CCL5, CUX2, FN1, MTSS1, NCF2, SEMA6A, SHANK3 |

**Supplementary Table 3:**
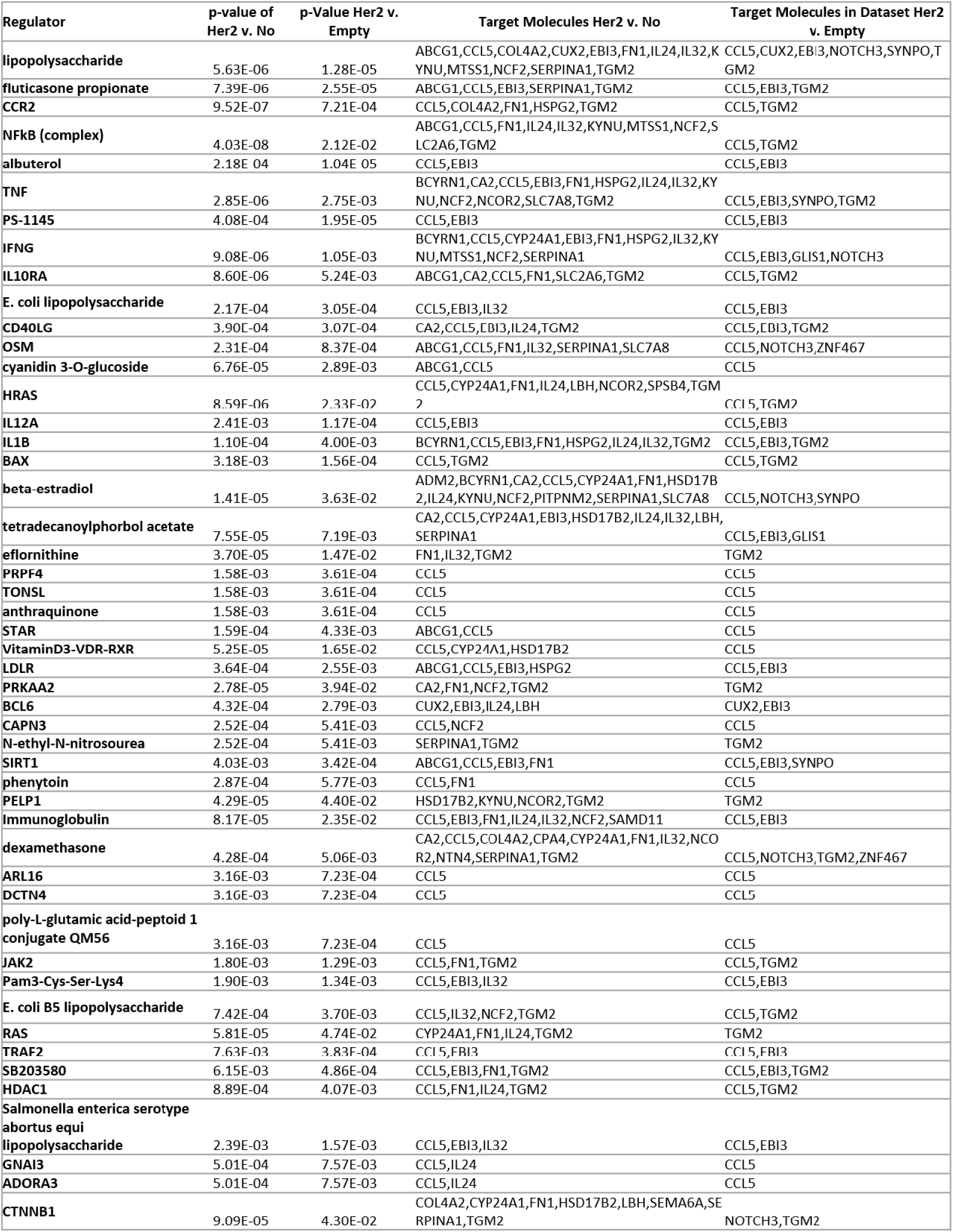
Upstream Factor analysis reveals potential regulators responding to Her2 treatment. Upstream regulators include mediators of inflammation such as LPS, TNF, IFNgamma, Fluticasone Propionate, tetradecanoylphorbol acetate, and/or dexamethasone.

## Notes

### Competing Interest Statement

The authors have declared no competing interest.

