## Supplementary material for "Nanolipoprotein mediated Her2 protein transfection induces malignant transformation in human breast acinar cultures": Suplemental figs

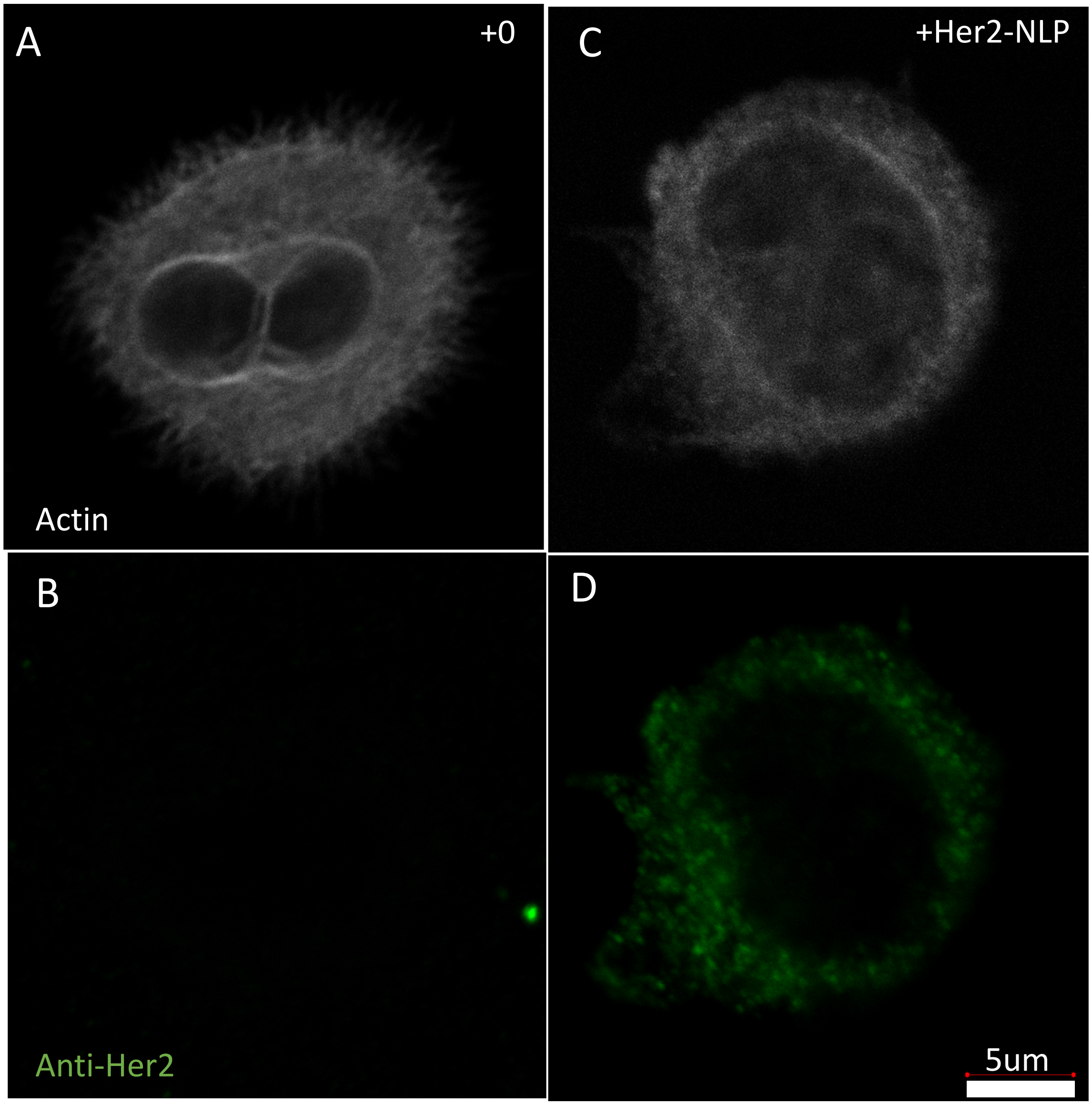
A
+0
Actin
C
+Her2-NLP
B
Anti-Her2
D
5um

**A.**

| Comparison | Up | Down | Total DEG |
|---|---|---|---|
| No_treatment-vs-empty_NLP | 0 | 1 | 1 |
| empty_NLP-vs-Her2_NLP | 9 | 0 | 9 |
| No_treatment-vs-Her2_NLP | 32 | 6 | 38 |

**B.**

| No Treatment v. Empty |
|---|
| BCYRN1 |
| **Empty v. Her2-NLP** |
| NOTCH3 |
| EBI3 |
| CUX2 |
| SYNPO |
| GLIS1 |
| ZNF467 |
| TGM2 |
| AC009133.2 |
| CCL5 |

| No Treatment v. Her2-NLP |
|---|
| IL32 |
| CYP24A1 |
| NTN4 |
| HSD17B2 |
| PITPNM2 |
| SLC7A8 |
| SEMA6A |
| CA2 |
| EBI3 |
| CUX2 |
| FN1 |
| KYNU |
| NCF2 |
| ADM2 |
| CPA4 |
| COL4A2 |
| HSPG2 |
| RNF175 |
| ABCG1 |
| SLC2A6 |
| IL24 |
| C15orf39 |
| RNF150 |
| MTSS1 |
| SPSB4 |
| SAMD11 |
| NCOR2 |
| SERPINA1 |
| TGM2 |
| RBM20 |
| LBH |
| LINC00863 |
| BCYRN1 |
| SHANK3 |
| AL132780.2 |
| IGFL2-AS1 |
| CCL5 |
| AL161431.1 |

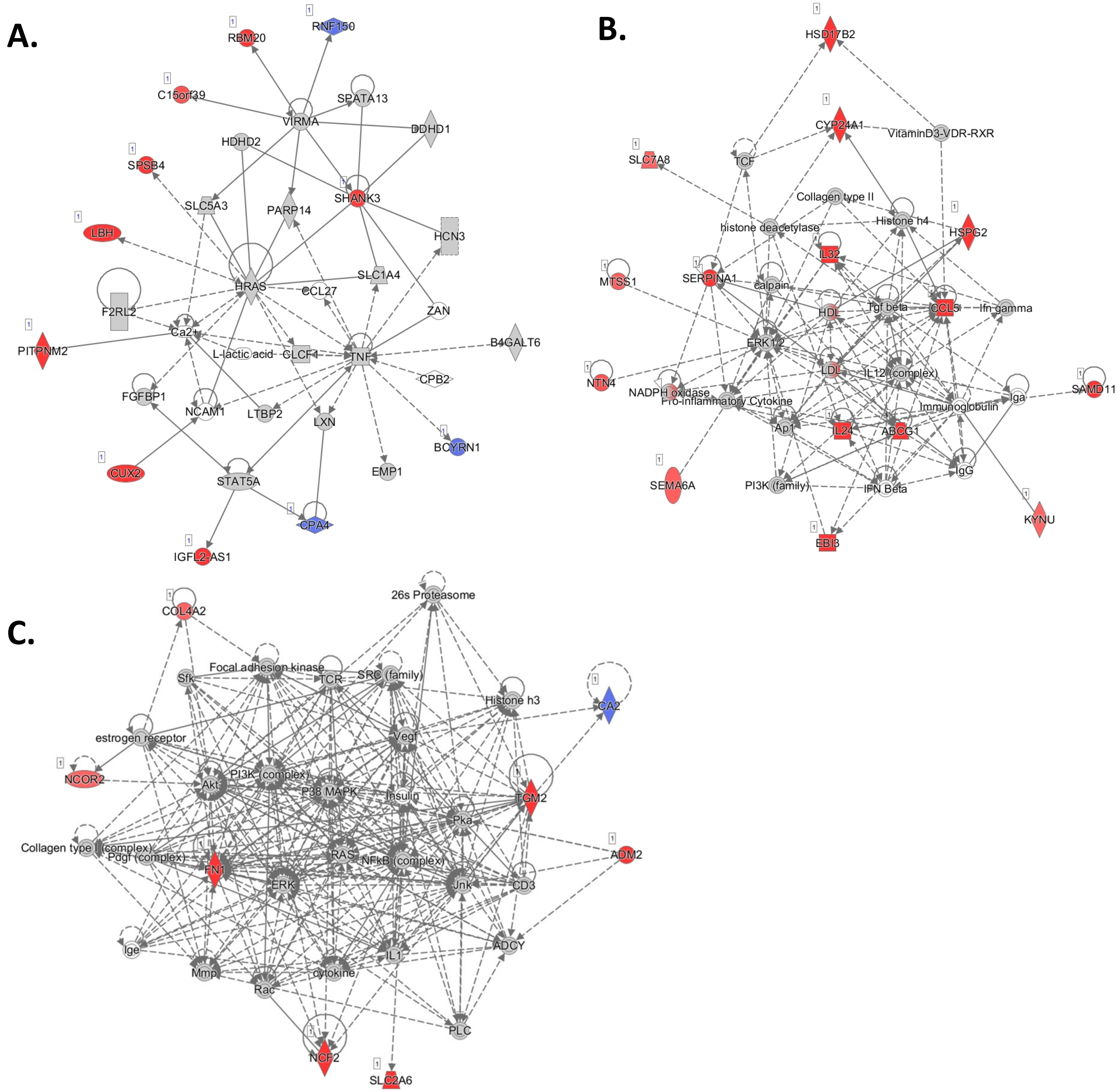
A.
RNF150
RBM20
C15orf39
SPATA13
VIRMA
DDHD1
HDHD2
SPSB4
SHANK3
SLC5A3
PARP14
LBH
HCN3
SLC1A4
HRAS
CCL27
F2RL2
ZAN
Ca2+
PITPNM2
B4GALT6
L-lactic acid
CLCF1
TNF
CPB2
FGFBP1
NCAM1
LTBP2
LXN
BCYRN1
EMP1
CUX2
STAT5A
CPA4
IGFL2-AS1
B.
HSD17B2
CYP24A1
VitaminD3-VDR-RXR
SLC7A8
TCF
Collagen type II
Histone h4
histone deacetylase
HSPG2
IL32
MTSS1
SERPINA1
calpain
HDL
Tgf beta
CCL5
Ifn gamma
ERK1/2
NTN4
LDL
IL12 (complex)
SAMD11
NADPH oxidase
Pro-inflammatory Cytokine
Iga
Immunoglobulin
Ap1
IL24
ABCG1
SEMA6A
PI3K (family)
IFN Beta
IgG
KYNU
EBI3
C.
26s Proteasome
COL4A2
Focal adhesion kinase
Sfk
TCR
SRC (family)
Histone h3
CA2
estrogen receptor
Vegf
NCOR2
PI3K (complex)
Akt
P38 MAPK
Insulin
TGM2
Pka
Collagen type I (complex)
Pdgf (complex)
RAS
NFkB (complex)
ADM2
FN1
ERK
Jnk
CD3
Ige
ADCY
IL1
Mmp
cytokine
Rac
PLC
NCF2
SLC2A6

**A.**

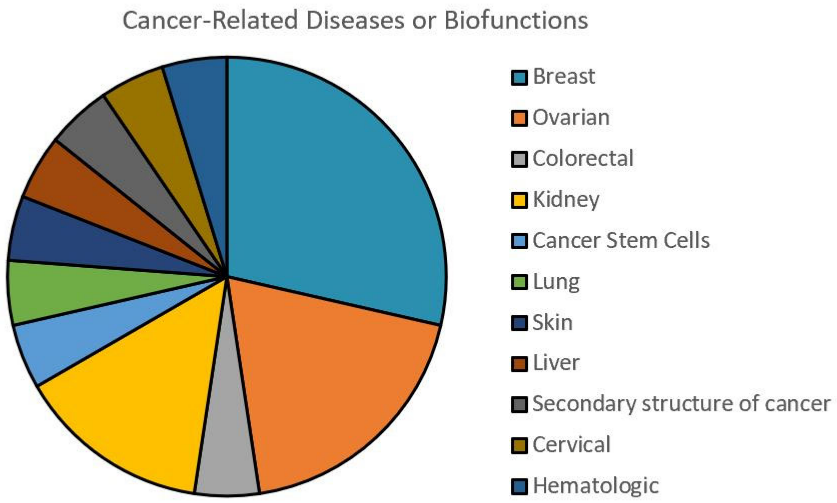


**B.**

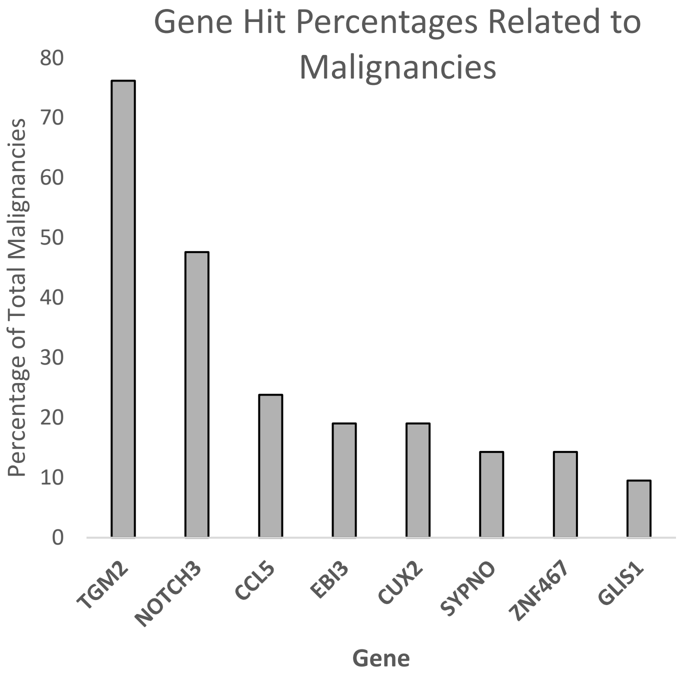


**C.**

| Cancer Type | Related malignancies | Molecular Hits | p-value |
|---|---|---|---|
| Breast | Migration of breast cancer cell lines | CCL5,EBI3,TGM2 | 0.000365 |
| | Attachment of breast cancer cell lines | TGM2 | 0.00211 |
| | Dissemination of breast cancer cell lines | CCL5 | 0.00457 |
| | Inflammatory breast cancer | NOTCH3 | 0.0143 |
| | Cell proliferation of breast cancer cell lines | NOTCH3,TGM2 | 0.0279 |
| | Primary breast cancer | NOTCH3 | 0.0292 |
| Ovarian | Modification of ovarian cancer cell lines | TGM2 | 0.000704 |
| | Epithelial-mesenchymal transition of ovarian cancer cell lines | TGM2 | 0.00141 |
| | Cell proliferation of ovarian cancer cell lines | NOTCH3,TGM2 | 0.00184 |
| | Sphere formation of ovarian cancer cell lines | TGM2 | 0.00211 |
| Kidney | Colony formation of kidney cancer cell lines | TGM2 | 0.00281 |
| | Apoptosis of kidney cancer cell lines | TGM2 | 0.0188 |
| | Proliferation of kidney cancer cell lines | TGM2 | 0.0364 |
| Cancer Stem Cells | Expansion of cancer stem cells | NOTCH3 | 0.00457 |
| Liver | Hepatobiliary system cancer | CUX2,GLIS1,NOTCH3,SYNPO,TGM2,ZNF467 | 0.0119 |
| Lung | Cell death of lung cancer cell lines | NOTCH3,TGM2 | 0.00853 |
| Cervical | Migration of cervical cancer cell lines | TGM2 | 0.0353 |
| Colorectal | Colorectal cancer | CCL5,CUX2,EBI3,NOTCH3,SYNPO,TGM2, ZNF467 | 0.00235 |
| Hematologic | Hematologic cancer | CCL5,CUX2,EBI3,NOTCH3 | 0.0472 |
| Secondary structure of cancer | Cancer of secretory structure | CCL5,CUX2,EBI3,GLIS1,NOTCH3,SYNPO,TGM2, ZNF467 | 0.0296 |
| Skin | Migration of skin cancer cell lines | TGM2 | 0.00946 |
